# Structure of the Leiomodin-2 Regulated Actin Filament Pointed End Assembly from Profilactin

**DOI:** 10.64898/2026.05.14.725239

**Authors:** Cristina M. Risi, Tania M. Larrinaga, Alla S. Kostyukova, Carol C. Gregorio, Vitold E. Galkin

**Author notes:** Corresponding author: Vitold E. Galkin.

## Abstract

Cardiac contraction depends on synchronized interactions between myosin-based thick filaments and actin-based thin filaments (TFs). Precise regulation of TFs length is vital for cardiac function, as any alteration in length leads to severe myopathies. Actin filaments form the backbone of the TF and have two unequal ends – fast-growing barbed and slow-growing pointed. In muscle, the barbed end is capped at the Z-line, while the pointed end is regulated by the tropomodulin family of proteins. Tropomodulin caps the pointed end, while leiomodin-2 (Lmod2) promotes actin nucleation and pointed end elongation. Lmod2 has a unique C-terminal extension (CTE) that is important for actin nucleation and binds to the sides of matured TFs. The structural mechanism by which Lmod2 promotes elongation remains elusive. We employed cryo-electron microscopy to visualize the structure of growing pointed ends nucleated by Lmod2 from profilactin. We show that Lmod2’s leucine-rich repeat domain (LRR) stabilizes terminal actin subunits by binding across the helical groove of actin. We identified two distinct populations of pointed-end LRR-containing complexes on one or both actin strands. LRR binding pushes the terminal actins outward from their ideal positions in the actin filament, introducing strain at the pointed end that squeezes LRR from the filament’s exterior. We also show that the Lmod2 CTE may stabilize Lmod2 binding to the pointed end. We suggest that Lmod2 promotes the addition of new actins to the pointed end but is expelled from the growing filament, thereby maintaining the concentration of Lmod2 required for further elongation.

## INTRODUCTION

In cardiac muscle, cyclical contraction and relaxation are produced by the interactions between myosin-containing thick filaments and actin-based thin filaments (TFs) arranged in a crystalline-like lattice within the muscle’s contractile units—the sarcomeres [1, 2]. Despite continuous protein turnover [3], precise regulation of the TF length is essential for the proper function of striated muscle, as it determines the degree of their overlap with thick filaments and thereby, according to the sliding filament model[4], controls force generation. Consequently, even modest perturbations in filament length are associated with severe skeletal and cardiac myopathies (reviewed in [5–7]). While early studies emphasized structural proteins such as nebulin as molecular rulers [8], it is now accepted that TFs’ length is dynamically regulated by a network of actin-associated proteins acting at the pointed filament ends – leiomodin (Lmod) and tropomodulin (Tmod). Tmod forms a tight cap at the pointed end to prevent actin polymerization and depolymerization [9, 10]. Lmod nucleates actin in vitro [11] and acts as a “leaky cap” at the TF pointed end to allow its elongation [12–14]. In cells, monomeric actin is complexed with the actin-sequestering protein profilin (profilactin), which prevents its uncontrolled polymerization under physiological salt conditions in the cytoplasm. In myocytes, profilin modulates sarcomeric organization via its actin sequestration activity, which is essential for maintaining the structural integrity of the sarcomere [15].

The Lmod family is structurally related to the Tmod family, sharing several conserved actin-binding sites (ABS) and a tropomyosin binding site (TmBS) [5, 7] (Fig. 1A and B). Instead of the second TmBS found in Tmod, Lmod2 has a linker between its ABS1 and leucine-rich repeat (LRR) domains that contains an acidic region (Fig. 1A, EDRR), which may regulate Lmod’s nucleation activity in a Ca^2+^-dependent manner [16]. The LRR domain has been shown to bind actin and promote actin nucleation [17]. However, unlike Tmods, Lmods possess a unique ∼150-residue C-terminal extension (CTE) that harbors three additional ABS [18, 19] (Fig. 1A cyan), one of which is a Wiskott-Aldrich homology 2 (Fig. 1A, WH2) domain that facilitates the recruitment of actin monomers to promote polymerization [20]. Deletion of the CTE greatly inhibits Lmod-dependent actin nucleation/polymerization [18]. Importantly, the poly-proline (poly-P) region located between the first two ABS of the CTE (Fig. 1A, poly-P) has been shown to bind profilin, which corroborates the ability of Lmod to effectively promote polymerization of profilactin [19]. Finally, the CTE has been shown to bind to the sides of fully matured cardiac TFs in a Ca^2+^-dependent manner, presumably to protect newly formed thin filament units during systole [18].

**Figure 1.**
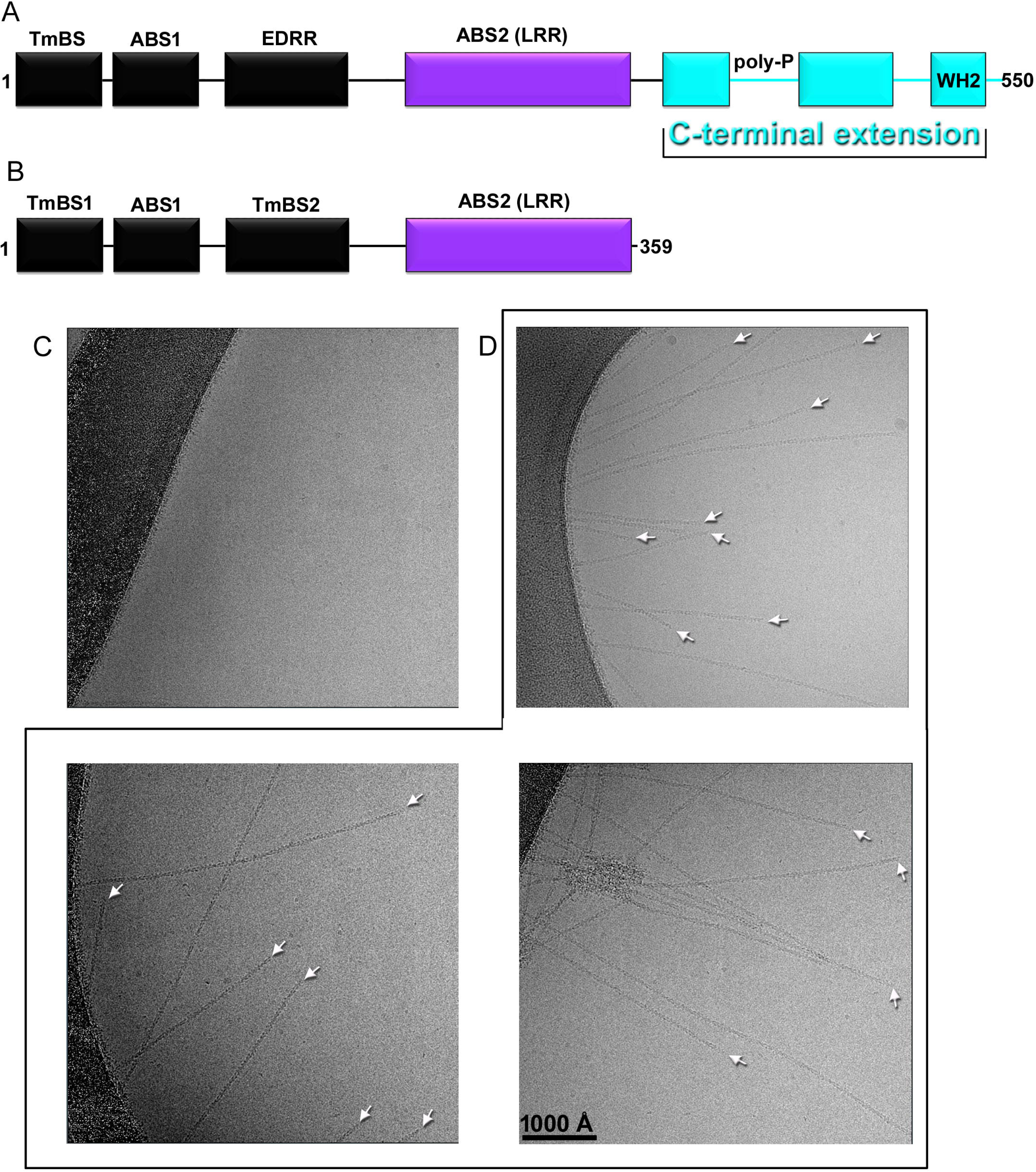
The domain organization of Lmod2. Lmod2 promotes profilactin polymerization. (A) Leiomodin 2 (Lmod2) consists of a tropomyosin-binding site (TmBS), an actin-binding site 1 (ABS1), and the glutamic acid/aspartic acid-rich region (EDRR), depicted as black rectangles, followed by the second actin-binding site (ABS2), known as the leucine-rich region (LRR, purple rectangle). The C-terminal extension (cyan) is unique to Lmod2 and comprises three actin-binding sites (cyan boxes), one of which is the WH2 domain, and the poly-P region, which binds profilin. (B) Tmod comprises TmBS1, ABS1, and TmBS2 (black rectangles), followed by the LRR domain (purple rectangle) that forms the second actin-binding site (ABS2). Tmod lacks a C-terminal extension. (C and D) Cryo-EM micrographs show that profilactin does not form filaments at high salt without Lmod2 (C) but readily polymerizes in the presence of Lmod2 (D). The ends of actin filaments are marked with white arrows. The scale bar is 1000 Å (0.1 µm).

All the abovementioned facts demonstrate the interplay of multiple functional sites of Lmod, which allow it to precisely control TF length in a multicomponent, highly dynamic, and mechanically stressed system such as the cardiac sarcomere. Nevertheless, the precise molecular mechanisms by which these functional sites promote actin polymerization at the pointed end are unknown. Here, we use cryo-electron microscopy (cryo-EM) to elucidate the structure of the actively growing pointed ends in the presence of profilactin. We show that Lmod2 coordinates two actin subunits at the pointed end of the actin filament on one or two actin strands through its LRR domain, while its CTE interacts with actin protomers at a distance from (below) the pointed end. The arrangement of LRR-bound actins differs from that of mature F-actin, indicating that once polymerization is complete, LRR is expelled from the actin filament and cannot stay bound to the sides of F-actin. Our data suggests that the interaction of the profilin-binding CTE with actin may not only promote LRR-dependent stabilization of the terminal actin protomers, but also foster the release of the profilin from the lmod2-profilin complex.

## RESULTS

To elucidate the mechanism of Lmod2-driven actin polymerization, we sought to determine the structure of the actively growing actin filament ends derived from the profilactin. In the absence of Lmod2, profilactin does not form filaments (Fig. 1C) because nucleation is impaired by profilin [21]. However, in the presence of Lmod2, actin-profilin readily forms filaments (Fig. 1D). In our conditions (1µM ATP-G-actin, 18 µM profilin, and 0.1 µM Lmod2) filaments were nucleated by Lmod2 [19], followed by their elongation from both ends with an estimated rate of elongation of ∼7 molecules/s at the barbed end [22] and ∼1 molecules/s at the pointed ends [23]. To maximize the number of filament ends suitable for image analysis (Fig. 1D, white arrows), we used the shortest possible time (∼45 to 60 sec) from mixing actin and profilin with Lmod2 to freezing the sample. Over this period, we expect that the barbed end could add from 315 to 360 actin protomers, while the pointed end could add from 45 to 60 new subunits, yielding filaments ranging from 1.0 to 1.2 µm in length (assuming axial rise of actin is 27.5 Å). The measured lengths of negatively stained actin filaments polymerized for 60 sec under the same conditions as in cryo-EM (Fig. S1) were consistent with these estimates. Consistent with this, the number of ends per cryo-EM image was relatively small, resulting in a modest number of segments used for image analysis (n=21,712). The resultant reconstructions of the filament ends are shown in Fig. 2A-D (gray transparent surfaces), along with the corresponding frequencies (Fig. 2A-D, black numbers). The reconstruction protocol is detailed in the Supplementary Information methods section and depicted in Figures S2-S4. As expected, we obtained approximately equal populations of barbed (∼57%) and pointed (∼43%) ends.

**Figure 2.**
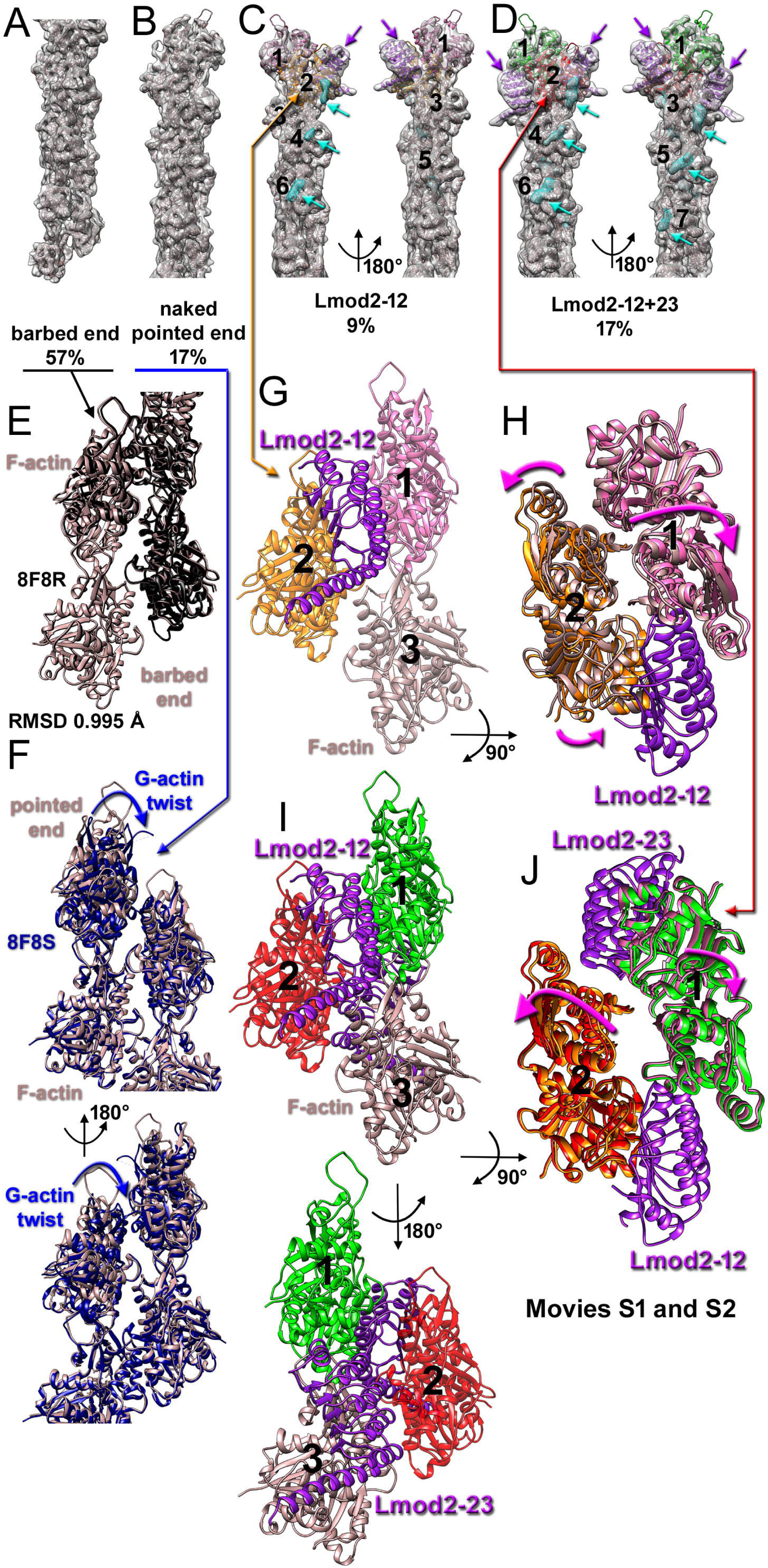
The structure of the barbed and pointed ends of actin filaments formed from profilactin in the presence of Lmod2. (A-D) The cryo-EM reconstructions (grey surface) and associated models (colored ribbons) of the barbed end (A), naked pointed end (B), pointed end with one Lmod2 bound (Lmod2-12), and pointed end with two Lmod2 molecules bound (Lmod2-12+23 (D). Two views related by the 180° azimuthal rotation are shown for pointed ends decorated with Lmod2 (C and D). The densities attributed to Lmod2 CTE are shown as a cyan mesh and marked with cyan arrows in C and D. Models have actin molecules in rosy-brown, except for the terminal actin protomers in C and D (pink, orange, green, and red), which possess altered conformations due to interactions with the LRR domain (purple ribbons and arrows). Actin protomers are numbered in black. The classes’ relative frequencies are indicated below the 3D reconstructions. (E) Barbed end structure from this study (rosy-brown ribbons) and structure of the barbed end of skeletal F-actin (PDB 8F8R) [24] (black ribbons) are nearly identical. (F) The naked pointed end from this study (rosy-brown ribbons) does not possess G-actin twist (blue arrows) in its terminal actin protomers, as observed in pointed ends of skeletal F-actin (PDB 8F8S)[24] (navy blue ribbons). (G) Model of Lmod2-12 side view. Position of Lmod2 LRR domain (purple ribbons) bound to actins 1 (pink ribbons) and 2 (orange ribbons). (H) Model of Lmod2-12 top view. LRR domain displaces actins 1 (pink ribbons) and 2 (orange ribbons) from their positions in the naked pointed end (rosy-brown ribbons) as indicated by magenta arrows. (I). Model of Lmod2-12+23 side view. Position of two Lmod2 LRR domains bound to actins 1 (green ribbons) and 2 (red ribbons), and to actins 2 (red ribbons) and 3 (rosy-brown ribbons). Two views related by a 180° azimuthal rotation are shown. (J) Model of Lmod2-12+23 top view. The second LRR domain displaces actins 1 (pink ribbons) and 2 (orange ribbons) from their positions in Lmod2-12 as indicated by magenta arrows. Movies S1 and S2 show the transition from the naked pointed end to Lmod2-12+23.

The 3-dimensional (3D) reconstructions of the barbed ends showed only actin density (Fig. 2A), and the atomic model derived from the 6 Å resolution map (Fig. S4B) was indistinguishable from the structure of the F-actin barbed end (PDB 8F8R) [24] with RMSD of 0.995 Å (Fig. 2E). Meanwhile, of ∼43% of segments assigned to the pointed end class, ∼17% were found to represent pointed ends without any density that could be attributed to Lmod2 (Fig. 2B). The evaluation of the 8Å resolution reconstruction of that class (Fig. S4D) revealed that, in contrast to the self-polymerizing F-actin pointed end structure (PDB 8F8S)[24], the two terminal actin subunits were not in the G-actin state – i.e., did not possess the so-called G-actin twist (Fig. 2F, blue arrows). For that reason, we retrofitted the map with the central part of the actin filament from 8F8S [24], which naturally represented the F-actin structure. All the actin subunits yielded a perfect fit, except minor rigid body refitting was required for the terminal top actin protomer. The resultant model yielded an RMSD of 0.493 Å with the central part of 8F8S (Fig. S5). Given that pointed ends could not grow from profilactin without Lmod2 assistance [23], we concluded that those pointed ends represented either pointed ends from which Lmod2 molecules had dissociated or central parts of actin filaments that had been mechanically broken upon cryo-EM sample preparation.

Among the pointed ends (43% of all ends), 26% displayed additional densities bound to their terminal actin subunits, with 9% showing one additional density (Fig. 2C, purple arrows) and 17% showing two densities (Fig. 2D, purple arrows). Double occupancy was more frequent (17%) than single occupancy (9%). Since a single-occupied pointed end has Lmod2 bound to actins 1 and 2 (Fig. 2C and G-H, black numbers), we refer to this structure as “Lmod2-12”. Because the double-occupied ends have one Lmod2 molecule bound to actins 1 and 2 and the second bound to actins 2 and 3 of the opposite strand (Fig. 2D and I-J, black numbers), we refer to this structure as “Lmod2-12+23”. The resolution estimations for these two classes were 8.5 Å (Fig. S4F) and 8.1 Å (Fig. S4H), respectively. The additional densities bound to the tips of these ends perfectly matched the structure of the actin-binding LRR domain of Lmod2 (PDB 5WFN) [17] (Fig. 2C and D, purple arrows and ribbons). The LRR domain structure was docked into the Lmod2-12 map with minimal perturbations in the C-terminal long α-helix (Fig. S6A) and yielded an RMSD of 0.745 Å. The same was true for the Lmod2-12+23 map, except that the long C-terminal α-helix had to be rotated by ∼15° from its position in the actin-Lmod2 crystal structure (PDB 5WFN) [17] (Fig. S6B), which increased the RMSD between the model and the crystal structure (PDB 5WFN) [17] to 2.318 Å. Due to the insufficient resolution, we could not determine the molecular interface between the LRR domain of Lmod2 and actins 1 and 2 or 2 and 3. Nevertheless, our structures unambiguously demonstrate that the LRR domain stabilizes the interaction between the two actins across the strand (e.g., between actin 1 and 2 or between actin 2 and 3), thus promoting the addition of actin monomers to the pointed end. Notably, despite the filaments being formed from profilactin, we were unable to observe any density resembling profilin in any of our four structural classes (Fig. 2A-D).

Importantly, we found that the positioning of the two terminal actins (marked as 1 and 2 in Fig. 2G-J) in both Lmod2-12 (Fig. 2G and H) and Lmod2-12+23 (Fig. 2I and J) complexes differed from that in the naked pointed end. In the Lmod2-12, the two distal actins were moved outwards to accommodate the LRR domain bound to the helical groove of actin, splitting the two strands of the actin filament (Fig. 2H, magenta arrows, movies S1 and 2). In the Lmod2-12+23 structure, the second LRR domain of Lmod2 bound to the opposite strand pushed the two terminal actins even further away from their positions in the actin helix (Fig. 2J, magenta arrows, movies S1 and 2). These findings unquestionably show that the LRR domain cannot bind to the groove of F-actin of the actin-based cardiac thin filament (cTF) due to insufficient opening of the F-actin helical groove in the matured filament. The lack of LRR binding to cTF actin groove has been reported [18]. Also, the displacement of the terminal actin protomers from their ideal positions within the actin helix strongly suggests that upon elongation, LRR domains are expelled from the filament due to the formation of energetically favorable contacts between the two actin strands. Consistently, we experimentally established that the LRR domain does not bind to actins 2 and 3 of the filament pointed end alone and requires the LRR domain bound on top of it to actins 1 and 2 (Fig. S3 and Supplementary methods).

In addition to the LRR-domain density, the three actin protomers on one strand (Fig. 2C and 3A, cyan mesh and arrows) or on both strands (Fig. 2D and 3B, cyan mesh and arrows) adjacent to the LRR domain showed well-defined additional densities. These densities were inconsistent with the profilin structure, but their elongated shape and localization on the surface of actin were consistent with the three actin binding sites of Lmod2 CTE on the sides of cTF (Fig. 3C and D) [18]. Since these densities were absent at the barbed end or at the “naked” pointed end, where the LRR domain was missing, and their resemblance to the footprint of the Lmod2 CTE on the cTF (Fig. 3D), we attributed them to the CTE of Lmod2 bound to the pointed filament end.

**Figure 3.**
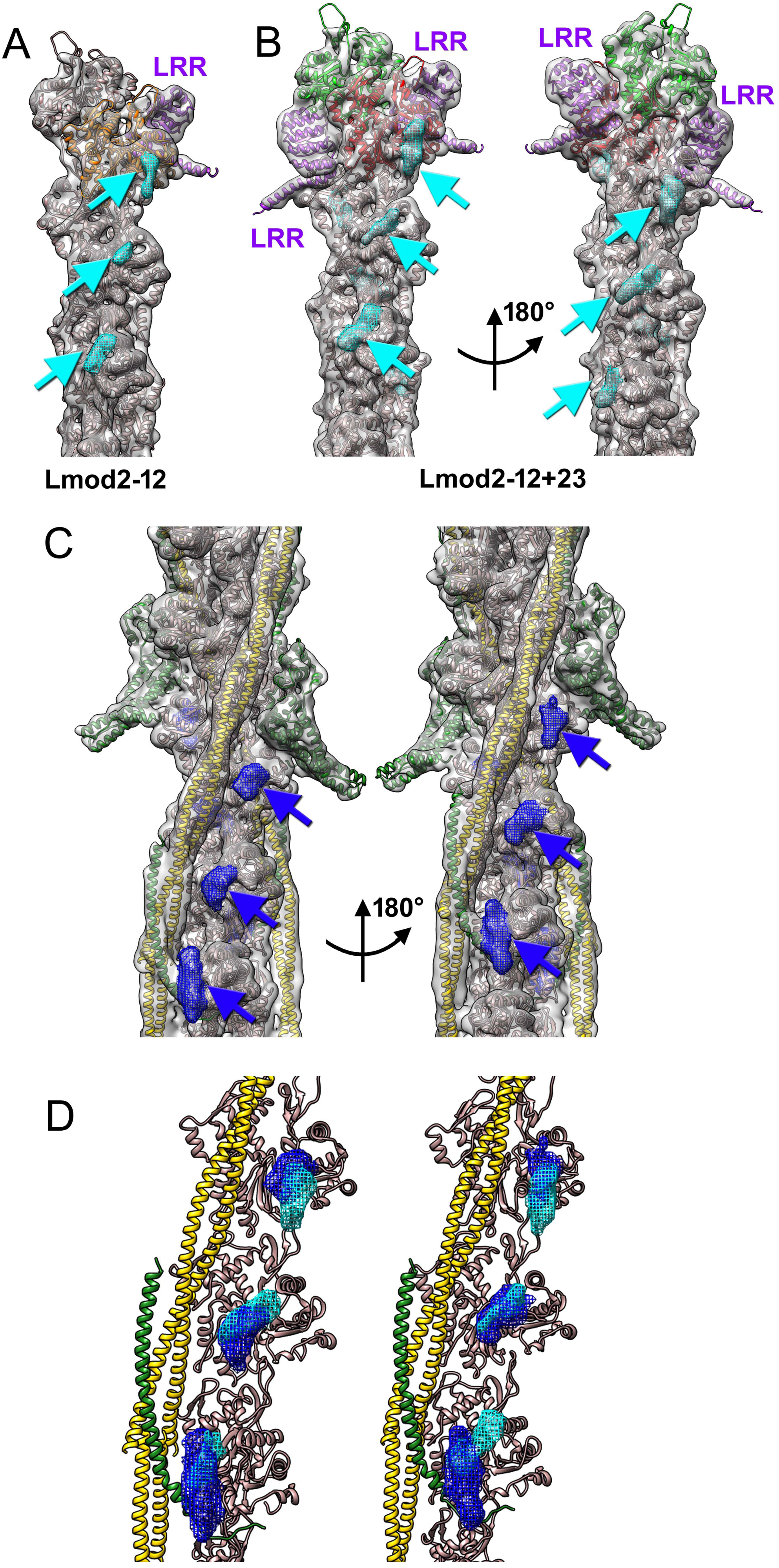
Comparison of Lmod2 CTE cTF side binding to side binding on the pointed filament end. (A and B) Density maps and corresponding models of the Lmod2-12 (A) and Lmod2-12+23 (B) classes. Additional density attributed to CTE is shown as a cyan mesh and indicated by cyan arrows. Ribbons are colored as in Fig. 1C and D. (C) Density map and corresponding model of Lmod2-decorated cTF from [18]. The CTE-attributed density is shown as a blue mesh and marked with blue arrows. Two views related by an azimuthal rotation of 180 ° are shown. Actin is rosy-brown ribbons, tropomyosin is gold ribbons, and the troponin complex and TnT1 are green ribbons. (D) Comparison of the size and positioning of the CTE densities found in the Lmod2-decorated cTFs (blue mesh) with the additional densities on the surface of actin of Lmod2-12 and Lmod2-12+23 classes (cyan mesh).

## DISCUSSION

Polymerization at actin filament pointed ends was long considered unfavorable under physiological conditions [25], especially when complexed with profilin [26]. However, recent studies identified at least two cellular factors that promote the addition of actin monomers to the pointed end – bacterial Vibrio toxins VopF and VopL [27] and mammalian muscle protein Lmod. Initial experiments implied that Lmod functioned as an actin nucleator [11]. However, subsequent studies suggested that Lmod could also promote actin assembly at the pointed end of the thin filament [5, 12–14, 28]. Finally, single-molecule experiments directly demonstrated Lmod’s elongation activity at the pointed ends of actin filaments [23]. However, the structural details of the elongating pointed ends remained elusive.

In contrast to Tmod (Fig. 1B), Lmod2 has an approximately 20-kDa C-terminal extension (Fig. 1A) that harbors a poly-P region and a WH2 domain (Fig. 1A, cyan rectangles). WH2 domains are well-known G-actin-binding motifs that bind actin monomers at the barbed end [20]. Notably, the WH2 domain of Lmod plays a fundamental role in actin nucleation [11, 17], while poly-P regions are found in actin-binding proteins that bind profilactin to promote G-actin polymerization [29]. Recently, we demonstrated that the poly-P region interacts with profilin, presumably facilitating profilactin polymerization at the pointed end [19]. Here, we analyzed the structure of the ends of actively growing actin filaments formed from profilactin.

In previous studies, a mixture of CapZ, Tmod, tropomyosin, and actin was used to produce short actin filaments, yielding more ends per micrograph[24]. Here, we aimed to study the structure of the actively growing ends to elucidate dynamic structural modes of Lmod2 interactions during elongation of pointed ends. The reduced number of available filaments’ ends reduced apparent resolution in the observed classes to 6 - 8Å (Fig. S4). However, the achieved resolution was sufficient to unambiguously dock the available high-resolution structures of actin (Fig. 2E and S5) and the LRR domain of Lmod2 (Fig. S6) into the experimental maps with minimal perturbations.

The length of actin filaments used for the image analysis (Fig S1) was consistent with the experimentally measured elongation rates from profilactin at the barbed end [22] and at the pointed end in the presence of Lmod2 [23]. Hence, under our experimental conditions, both ends of the actin filaments were actively growing. The structure of the barbed ends observed in this study fully agreed with the known high-resolution structure of the barbed end [24]. Nonetheless, the structure of the naked (i.e., without Lmod2) pointed ends in our study was drastically different from that of the pointed ends formed by self-polymerized G-actin [24] (Fig. 2F), suggesting that the mechanism of elongation at the pointed end was different, as expected, since profilactin cannot polymerize from the pointed end [22]. Most of the pointed ends possessed well-defined Lmod2 LRR-domain density attached to one (Fig. 2C) or both (Fig. 2D) actin strands. The most prominent finding when examining these two structures was the splitting of the actin filament’s two strands at its pointed end (Fig. 2H and J, Movies S1 and 2). The extent of splitting was larger when two LRR-domains were bound to the pointed end (Movie S2). Hence, while the LRR domain of Lmod2 promotes the attachment of new actin molecules to the pointed end by connecting them to existing actin protomers across the strand (e.g., Lmod2-12 connects actins 1 and 2, while Lmod2-12+23 connects actins 1 and 2, along with actins 2 and 3, respectively), it alters the lateral interactions between actin protomers that hold the two strands together (Movie S3). This supports the hypothetical processive mechanism for pointed-end elongation by Lmod2, as depicted in Figure 4: LRR domains that bind to actins 2 and 3 are expelled from the actin filament interior due to interactions between the two actin strands, allowing Lmod2 molecules to participate in new rounds of elongation rather than being trapped and bound to the body of the actin filament. The proposed mechanism is consistent with two of our findings: (1) we did not detect LRR domains bound to actins 2 and 3 alone or any traces of LRR domains bound below actin 3 (Fig. S2 and S3); and (2) LRR-domain binding to the cTF actin backbone was not found [18].

**Figure 4.**
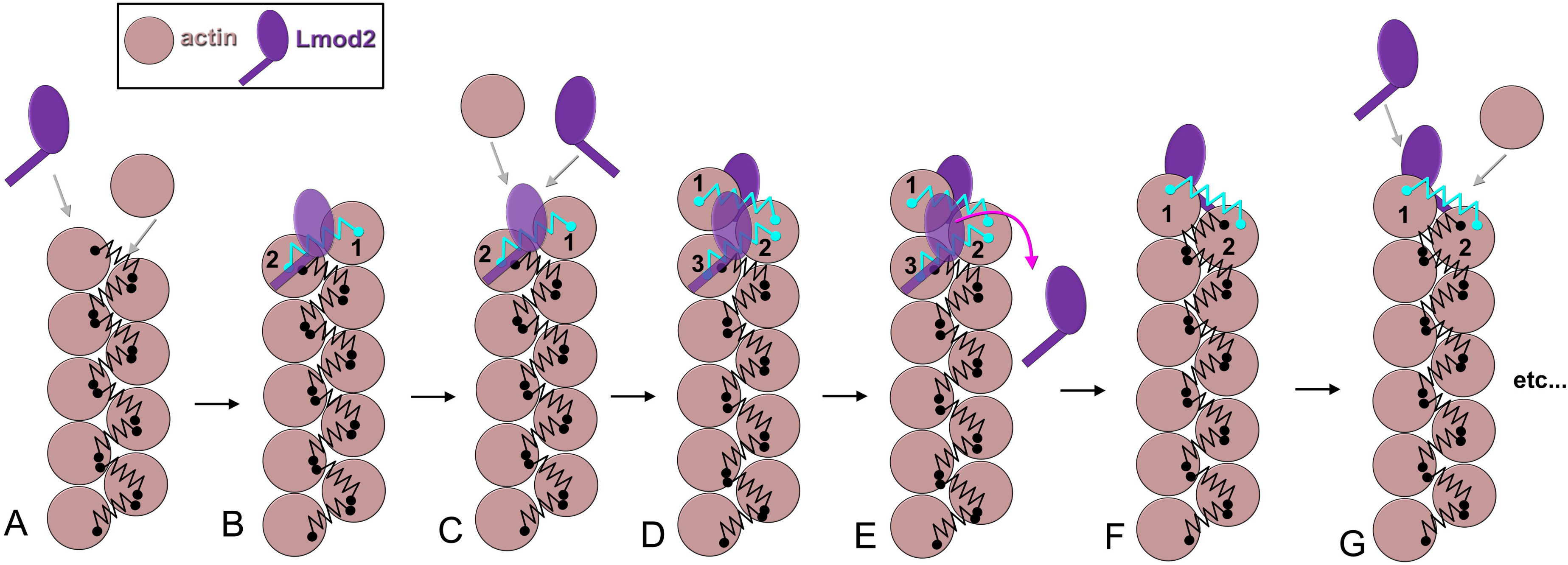
Hypothetical mechanism of pointed-end elongation by Lmod2. (A-G) Actin molecules are depicted as rosy-brown circles, Lmod2 molecules are in purple, and normal interstrand interactions between actin protomers that hold the two strands are shown as black springs. (A) Actin (complexed with profilin) and Lmod2 interact with the pointed end (gray arrows). (B) The LRR domain of Lmod2 cross-links the two terminal actins (actins 1 and 2) – one part of the original filament (actin 2), the other a new protomer (actin 1). The LRR domain splits the actin strands (interstrand interactions are shown as cyan springs) on actins 1 and 2. (C) Subsequent actin (complexed with profilin) and Lmod2 interact with the pointed end (gray arrows). (D) The new actin is added to the pointed end (actin 1), allowing the new LRR domain to cross-link actins 1 and 2, while the originally attached LRR remains on actins 2 and 3. The LRR domain (?) cross-linking actins 1 and 2 and the LRR domain connecting actins 2 and 3 enhance the splitting of the two actin strands (two cyan springs). (E) The tension caused by the split of the actin strands squeezes the LRR domain bound to actins 2 and 3 off (magenta curved arrow). (F) The pointed end with a single bound Lmod2 is ready to add the next actin. (G) The elongation process continues as a new G-actin molecule, along with Lmod2, approaches the growing pointed end.

In our previous work, we demonstrated that the CTE of Lmod2 is involved in interactions with the cTF actin backbone [18, 19]. In contrast to G-actin binding, only the C-terminal region of the WH2 domain participates in binding to the sides of cTF [19]. Notably, we observe a very similar pattern of interactions on the three actin protomers adjacent to the LRR domain (Fig. 3), which strongly suggests that the CTE of Lmod2 stabilizes Lmod2 attachment to the pointed end. The importance of Lmod2 CTE in keeping Lmod2 bound to the elongating pointed end has been directly demonstrated by the single-molecule techniques [23]. Also, it is feasible to speculate that CTE binding to actin protomers may help strip off profilin from profilactin, if Lmod2 binds to the pointed end complexed with profilactin via the WH2 domain. This hypothesis required further experimental work.

## MATERIALS AND METHODS

### Proteins and Buffers

Skeletal G-actin was purified as described in [30]. Full-length Lmod2 was obtained as described in [31]. G-buffer: 2 mM TRIS-HCl pH 7.8, 0.2 mM CaCl2, 0.2 mM ATP, 0.2 mM DTT. P-buffer: 20 mM TRIS-HCl pH 7.5, 150 mM KCl, 2 mM MgCl2, 2 mM EGTA, 1 mM DTT, 0.2 mM ATP.

### Cryo-EM

2 µM G-actin and 36 µM profilin were mixed in G-buffer. At the same time, 16.5 µM Lmod2 was diluted to 0.2 µM in P-buffer. 5 µl of each sample (1:1) were mixed to achieve final concentrations of 1 µM G-actin, 18 µM profilin, and 0.1 µM Lmod2 in a tube at room temperature for 30 sec. The mixture was applied to a glow-discharged lacey carbon grid at 15°C and 100% humidity in the Vitrobot Mark IV (FEI, Inc.) chamber. The sample was blotted with Whatman Grade 1 Filter Paper for 3 to 5 s and vitrified. A summary of imaging conditions and image reconstruction is provided in Supplementary Table S1. Image analysis was performed using RELION [32], while modeling was done using UCSF Chimera [33] and PHENIX software suites [34]. Experimental details are provided in Supplementary Materials and Methods.

### Measurement of filament length by negative staining TEM

The sample preparation was done as described for cryo-EM, except the aliquots were removed after 45 sec of incubation at room temperature and used for negative-stain EM. 7 μL of the sample was applied to a carbon-coated, glow-discharged grid (Electron Microscopy Sciences, FCF300-Cu-SB) and incubated for 15 sec before blotting with Whatman filter paper. Grids were then stained with 2% (w/v) uranyl acetate and imaged on a JEM-1200EX (JEOL) transmission electron microscope operated at 70 kV. Micrographs were recorded at a nominal magnification of 15,000X.

## Accession numbers

The cryo-EM maps have been deposited to the Electron Microscopy Data Bank (https://www.ebi.ac.uk/pdbe/emdb) with accession codes: 77128 for the barbed end, 77130 for the undecorated pointed end, 77131 for the pointed end with one Lmod2 bound (Lmod2-12), and 77132 for the pointed end with two Lmod2 bound (Lmod2-12+23).

## Acknowledgments

The study was supported by NIH grant R01 GM120137 (to VEG, CCG and ASK), NIH grant R01 HL73431 (to CCG). The authors thank Mason Summers for assistance in establishing nucleation conditions and Dr. Mert Colpan for providing profilin.

## REFERENCES

1. Huxley, H. E. (1972). Structural changes in the actin- and myosin-containing filaments during contraction. Cold Spring Harb. Symp. Quant. Biol. 37, 361–376.

2. Spudich, J. A., Huxley, H. E. & Finch, J. T. (1972). Regulation of skeletal muscle contraction. II. Structural studies of the interaction of the tropomyosin-troponin complex with actin. J. Mol. Biol. 72, 619–632.

3. Michele, D. E., Albayya, F. P. & Metzger, J. M. (1999). Thin filament protein dynamics in fully differentiated adult cardiac myocytes: toward a model of sarcomere maintenance. J Cell Biol. 145, 1483–1495. 10.1083/jcb.145.7.1483.

4. Huxley, H. & Hanson, J. (1954). Changes in the Cross-Striations of Muscle during Contraction and Stretch and their Structural Interpretation. Nature. 173, 973–976. 10.1038/173973a0.

5. Tolkatchev, D., Gregorio, C. C. & Kostyukova, A. S. (2022). The role of leiomodin in actin dynamics: a new road or a secret gate. FEBS J. 289, 6119–6131. 10.1111/febs.16128.

6. Iwanski, J., Gregorio, C. C. & Colpan, M. (2021). Redefining actin dynamics of the pointed-end complex in striated muscle. Trends Cell Biol. 31, 708–711. 10.1016/j.tcb.2021.06.006.

7. Fowler, V. M. & Dominguez, R. (2017). Tropomodulins and Leiomodins: Actin Pointed End Caps and Nucleators in Muscles. Biophys J. 112, 1742–1760. 10.1016/j.bpj.2017.03.034.

8. Labeit, S., Gibson, T., Lakey, A., Leonard, K., Zeviani, M., Knight, P., Wardale, J. & Trinick, J. (1991). Evidence that nebulin is a protein-ruler in muscle thin filaments. FEBS Letters. 282, 313–316.

9. Weber, A., Pennise, C. R., Babcock, G. G. & Fowler, V. M. (1994). Tropomodulin caps the pointed ends of actin filaments. J Cell Biol. 127, 1627–1635. 10.1083/jcb.127.6.1627.

10. Gregorio, C. C., Weber, A., Bondad, M., Pennise, C. R. & Fowler, V. M. (1995). Requirement of pointed-end capping by tropomodulin to maintain actin filament length in embryonic chick cardiac myocytes. Nature. 377, 83–86. 10.1038/377083a0.

11. Chereau, D., Boczkowska, M., Skwarek-Maruszewska, A., Fujiwara, I., Hayes, D. B., Rebowski, G., Lappalainen, P., Pollard, T. D. & Dominguez, R. (2008). Leiomodin is an actin filament nucleator in muscle cells. Science. 320, 239–243. 10.1126/science.1155313.

12. Tolkatchev, D., Smith, G. E., Jr., Schultz, L. E., Colpan, M., Helms, G. L., Cort, J. R., Gregorio, C. C. & Kostyukova, A. S. (2020). Leiomodin creates a leaky cap at the pointed end of actin-thin filaments. PLoS Biol. 18, e3000848. 10.1371/journal.pbio.3000848.

13. Larrinaga, T. M., Smith, G. E., Jr., Tolkatchev, D., Rast, T. J., Bunch, T. A., Colson, B. A., Pappas, C. T., Kostyukova, A. S. & Gregorio, C. C. (2026). N-Terminal Actin-Binding Site of Lmod2 Promotes Controlled Pointed End Elongation. Circ Res. 138, e327013. 10.1161/CIRCRESAHA.125.327013.

14. Mi-Mi, L., Farman, G. P., Mayfield, R. M., Strom, J., Chu, M., Pappas, C. T. & Gregorio, C. C. (2020). In vivo elongation of thin filaments results in heart failure. PLoS One. 15, e0226138. 10.1371/journal.pone.0226138.

15. Kooij, V., Viswanathan, M. C., Lee, D. I., Rainer, P. P., Schmidt, W., Kronert, W. A., Harding, S. E., Kass, D. A., Bernstein, S. I., Van Eyk, J. E. & Cammarato, A. (2016). Profilin modulates sarcomeric organization and mediates cardiomyocyte hypertrophy. Cardiovasc Res. 110, 238–248. 10.1093/cvr/cvw050.

16. Smith, G. E., Tolkatchev, D., Risi, C., Little, M., Gregorio, C. C., Galkin, V. E. & Kostyukova, A. S. (2022). Ca(2+) attenuates nucleation activity of leiomodin. Protein Sci. 31, e4358. 10.1002/pro.4358.

17. Boczkowska, M., Yurtsever, Z., Rebowski, G., Eck, M. J. & Dominguez, R. (2017). Crystal Structure of Leiomodin 2 in Complex with Actin: A Structural and Functional Reexamination. Biophys J. 113, 889–899. 10.1016/j.bpj.2017.07.007.

18. Little, M., Risi, C. M., Larrinaga, T. M., Summers, M. D., Nguyen, T., Smith, G. E., Jr., Atherton, J., Gregorio, C. C., Kostyukova, A. S. & Galkin, V. E. (2025). Interaction of cardiac leiomodin with the native cardiac thin filament. PLoS Biol. 23, e3003027. 10.1371/journal.pbio.3003027.

19. Summers, M. D., Little, M., Young, R. P., Osegueda, B., Palma Guillen, A., Smith, G. E., Jr., Cort, J. R., Galkin, V. E. & Kostyukova, A. S. (2026). Localization and functional exploration of leiomodin-2’s C-terminal binding sites. Biochim Biophys Acta Proteins Proteom. 1874, 141137. 10.1016/j.bbapap.2026.141137.

20. Dominguez, R. (2007). The beta-thymosin/WH2 fold: multifunctionality and structure. Ann N Y Acad Sci. 1112, 86–94. 10.1196/annals.1415.011.

21. Carlsson, L., Nystrom, L. E., Sundkvist, I., Markey, F. & Lindberg, U. (1977). Actin polymerizability is influenced by profilin, a low molecular weight protein in non-muscle cells. J Mol Biol. 115, 465–483. 10.1016/0022-2836(77)90166-8.

22. Courtemanche, N. & Pollard, T. D. (2013). Interaction of profilin with the barbed end of actin filaments. Biochemistry. 52, 6456–6466. 10.1021/bi400682n.

23. Biswas, S., Larrinaga, T., Choubey, S., Gregorio, C. C., Shekhar, S. (2026). Leiomodin 2 functions as a processive pointed-end elongator of actin filaments. bioRxiv. doi: 10.64898/2026.05.04.720848

24. Carman, P. J., Barrie, K. R., Rebowski, G. & Dominguez, R. (2023). Structures of the free and capped ends of the actin filament. Science. 380, 1287–1292. 10.1126/science.adg6812.

25. Pollard, T. D. (1986). Rate constants for the reactions of ATP- and ADP-actin with the ends of actin filaments. J. Cell Biol. 103, 2747–2754.

26. Pollard, T. D. & Cooper, J. A. (1984). Quantitative analysis of the effect of Acanthamoeba profilin on actin filament nucleation and elongation. Biochemistry. 23, 6631–6641.

27. Kudryashova, E., Ankita, Ulrichs, H., Shekhar, S. & Kudryashov, D. S. (2022). Pointed-end processive elongation of actin filaments by Vibrio effectors VopF and VopL. Sci Adv. 8, eadc9239. 10.1126/sciadv.adc9239.

28. Tsukada, T., Pappas, C. T., Moroz, N., Antin, P. B., Kostyukova, A. S. & Gregorio, C. C. (2010). Leiomodin-2 is an antagonist of tropomodulin-1 at the pointed end of the thin filaments in cardiac muscle. J Cell Sci. 123, 3136–3145. 10.1242/jcs.071837.

29. Ferron, F., Rebowski, G., Lee, S. H. & Dominguez, R. (2007). Structural basis for the recruitment of profilin-actin complexes during filament elongation by Ena/VASP. EMBO J. 26, 4597–4606. 10.1038/sj.emboj.7601874.

30. Ly, T., Moroz, N., Pappas, C. T., Novak, S. M., Tolkatchev, D., Wooldridge, D., Mayfield, R. M., Helms, G., Gregorio, C. C. & Kostyukova, A. S. (2016). The N-terminal tropomyosin- and actin-binding sites are important for leiomodin 2’s function. Mol Biol Cell. 27, 2565–2575. 10.1091/mbc.E16-03-0200.

31. Ly, T., Pappas, C. T., Johnson, D., Schlecht, W., Colpan, M., Galkin, V. E., Gregorio, C. C., Dong, W. J. & Kostyukova, A. S. (2019). Effects of cardiomyopathy-linked mutations K15N and R21H in tropomyosin on thin-filament regulation and pointed-end dynamics. Mol Biol Cell. 30, 268–281. 10.1091/mbc.E18-06-0406.

32. Scheres, S. H. (2012). RELION: implementation of a Bayesian approach to cryo-EM structure determination. J Struct Biol. 180, 519–530. 10.1016/j.jsb.2012.09.006.

33. Pettersen, E. F., Goddard, T. D., Huang, C. C., Couch, G. S., Greenblatt, D. M., Meng, E. C. & Ferrin, T. E. (2004). UCSF Chimera--a visualization system for exploratory research and analysis. J. Comput. Chem. 25, 1605–1612.

34. Liebschner, D., Afonine, P. V., Baker, M. L., Bunkoczi, G., Chen, V. B., Croll, T. I., Hintze, B., Hung, L. W., Jain, S., McCoy, A. J., Moriarty, N. W., Oeffner, R. D., Poon, B. K., Prisant, M. G., Read, R. J., Richardson, J. S., Richardson, D. C., Sammito, M. D., Sobolev, O. V., Stockwell, D. H., Terwilliger, T. C., Urzhumtsev, A. G., Videau, L. L., Williams, C. J. & Adams, P. D. (2019). Macromolecular structure determination using X-rays, neutrons and electrons: recent developments in Phenix. Acta Crystallogr D Struct Biol. 75, 861–877. 10.1107/S2059798319011471.

