## Supplementary Information for "Structure of the Leiomodin-2 Regulated Actin Filament Pointed End Assembly from Profilactin"

*Corresponding author: Vitold E. Galkin

**This PDF file includes:**

Supplementary text

Figures S1 to S6

Movie S1-S3

Tables S1

SI References

**Supplementary Information Text**

**Methods.**

**Imaging and image analysis.** Details of imaging conditions and image reconstruction are provided in Table S1. Micrographs (n=11,689) were collected on a 300-kV Titan Krios electron microscope equipped with a K3 direct electron detector and an energy filter operating in super-resolution mode. Micrographs were recorded using 40 subframes at a dose rate of ∼0.85 e-/Å2 per frame over a defocus range of 0.5–3.5 µm, with a pixel size of 0.678 Å. All processing was performed in RELION^1^. Images were imported into RELION for motion correction and dose weighting using internal RELION algorithms. The images were binned to a pixel size of 1.356 Å, and contrast transfer function (CTF) parameters were determined using CTFFIND4^2^. 21,712 particles representing filament ends were manually selected (Figure 2D, white arrows and Figure S2A, green circles). Particles were decimated to a raster of 2.712 Å/px (Figure S2B) and used for an initial classification by the polarity (i.e., barbed or pointed) of the filament end using models of the barbed and pointed ends generated from the available high-resolution structures (8F8R – barbed end; 5JLF – pointed end) filtered to 20 Å resolution (Figure S2C). We did not use the pointed end structure possessing G-twist of the terminal actin subunits (PDB 8F8S) to eliminate any bias in the sorting outcome. The resulting class average for the pointed end differed from the initial models due to the presence of LRR density (Figure S2C, purple arrows), while the barbed end class average was similar to the initial model. Overall, the sorting outcome was different from the initial references. Segments assigned to the barbed end (n=12,335) were used for 3D refinement (Figure S2D). Since the LRR-like density in the pointed end class average was smaller than expected, while the modes of binding were unknown, we used the particles assigned to the pointed end class (n=9,377) (Figure S2C, red box) for an unsupervised 3D classification into four classes (Figure S2E). Of note, LRR-domain presence indicated a full-sized Lmod2 molecule bound to the filament end; hence, we refer to it as the Lmod2-bound pointed end. Particles were re-extracted at a raster of 1.356 Å/px. The pointed-end model (Figure S2C) was filtered to 15 Å and used as the reference and mask (Figure S2E: white surface and grey mesh, respectively). The resulting class averages from the 4-class sorting are shown in Figure S2E. Class 1 (slate grey surface) had LRR-domain density on one strand (Figure S2E, purple arrow), class 2 (dark grey surface) represented pointed end without any LRR-domain density (naked pointed end), class 3 (light grey surface), along with class 4 (medium grey surface), possessed LRR-domain densities attached to both F-actin strands (Figure S2E, purple arrows). Hence, the unsupervised classification revealed three distinct classes of pointed ends: naked, pointed end with one Lmod2 molecule bound to actins 1 and 2 (e.g., Lmod2-12), and pointed end with two Lmod2 molecules bound – one to actins 1 and 2 and the other to actins 2 and 3 (e.g., Lmod2-12+23). Based on the findings from the unsupervised classification, we generated three references: naked pointed end, pointed end with Lmod2-12, and pointed end with Lmod2-12+23 (Figure S2F). The resulting class averages and corresponding frequencies from the 3-class sorting are shown in Figure S2G: naked class (n=3,762) (medium grey surface), Lmod2-12 class (n=1,992) (light grey surface), and Lmod2-12+23 class (n=3,623) (dark grey surface). To check if Lmod2 may bind to actins 2 and 3 (e.g., Lmod2-23), we generated a fourth reference (Figure S3A, dark grey surface) and used it along with the three models from the supervised sorting (Figure S2F). The resultant classes confirmed previously detected modes of binding (e.g., naked, Lmod2-12, and Lmod2-12+23), while the fourth class represented a partially misaligned Lmod2-12 class (Figure S3C).

The particles representing the barbed end class (Figure S2E, dark grey surface) were re-extracted, refined, and post-processed at 1.356 Å/px (Figure S4A), using the 3D class-average low-pass filtered to 15 Å as the starting reference and mask (Figure S4A, red surface and grey mesh, respectively). The density map was segmented (Figure S4A, cerulean surface) in USCF Chimera^3^ using the segmentation tool^4^ and used for modeling (Figure S4A, rosy-brown ribbons). The global resolution was calculated using RELION internal algorithms^1^ using 0.143 FSC criterion (Figure S4B).

The naked pointed end class particles (Figure S2H, medium grey surface) were re-extracted, refined, and post-processed at 1.356 Å/px (Figure S4C), using the 3D class-average low-pass filtered to 15 Å as the starting reference and mask (Figure S4C, green surface and grey mesh, respectively). The density map was segmented (Figure S4C, yellow surface) in USCF Chimera^3^ using the segmentation tool^4^ and used for modeling (Figure S4C, rosy-brown ribbons). The global resolution was calculated using RELION internal algorithms^1^ using 0.143 FSC criterion (Figure S4D).

Particles having Lmod2 bound to actins 1 and 2 (e.g., Lmod2-12) (Figure S2H, light grey surface) were processed in the same manner. The 1.356 Å/px re-extracted particles were used for 3D refinement, with the 3D class-average low-pass filtered to 15 Å as the starting reference and mask (Figure S4E, pink surface and grey mesh, respectively), and post-processed (Figure S2E). The density map was segmented (Figure S4C, lavender surface) and used for modeling (Figure S4E, colored ribbons; using the same color code as in Figure 2). The global resolution was calculated using RELION's internal algorithms^1^ using 0.143 FSC criterion (Figure S4F).

Particles having two Lmod2 molecules bound (e.g., Lmod2-12+23) (Figure S2H, dark grey surface) were processed in the same manner as previously described. The 3D class-average low-pass filtered to 15 Å was used as the starting reference and mask (Figure S4G, cyan surface and grey mesh, respectively), post-processed, segmented (Figure S4G, magenta surface), and used for modeling (Figure S4G, colored ribbons: using the same color code for the model as Figure 2. The global resolution was calculated using RELION internal algorithms^1^ using 0.143 FSC criterion (Figure S4H).

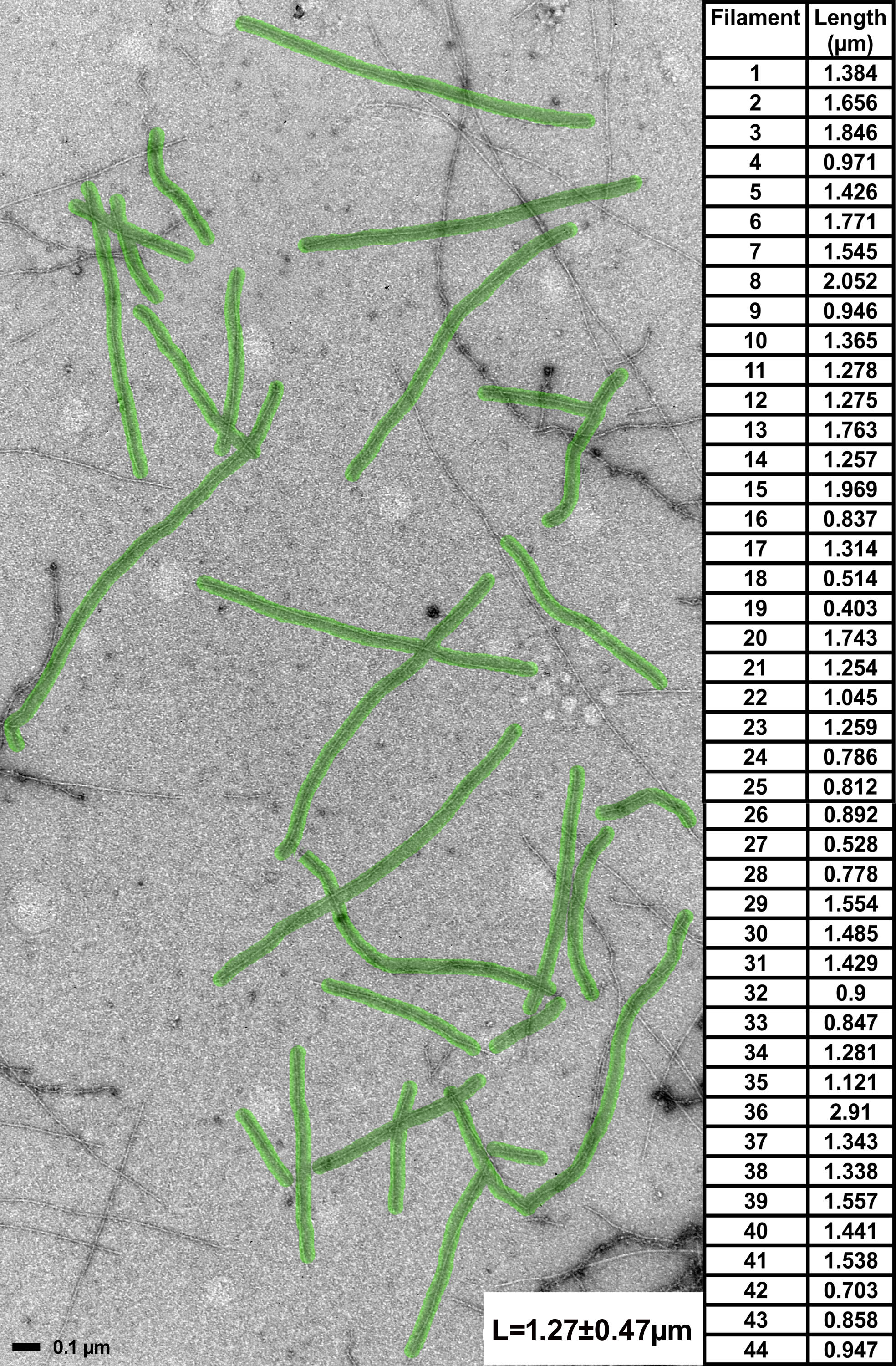

**Figure S1. Length estimation of actin filaments formed from profilactin in the presence of Lmod2.** An example of a TEM image of negatively stained actin filaments has filaments used for length measurement marked in transparent green. The experimental values (n=44) are shown in the table on the right, along with the mean and its standard deviation.

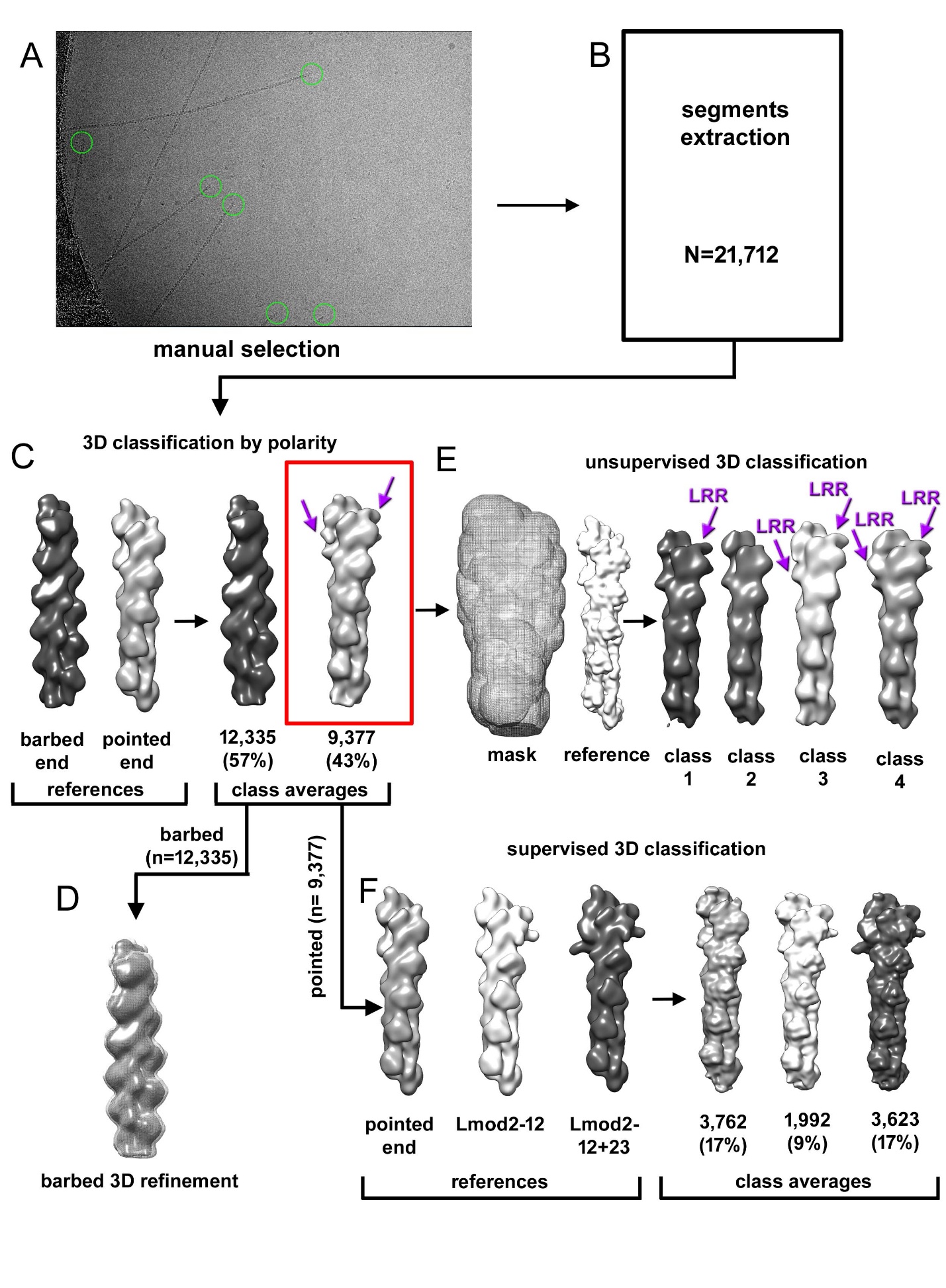

**Figure S2. 3D classification algorithm for filament ends.** (A-B) Filament ends were manually selected, as indicated by green circles, and extracted from cryo-EM micrographs. (C) In the first round, particles were sorted by the polarity of their ends. References and class averages are shown with corresponding frequencies. (D) The particles assigned to the barbed end were used for 3D refinement, with the class average as the mask and reference, as detailed in Figure S3A. (E) The naked, pointed-end class average was filtered to 15 Å to be used as a reference and a mask (white surface and grey mesh, respectively) for the unsupervised sorting of the pointed-end particles. The resultant class averages for the four classes are shown: class 1 (slate grey surface), class 2 (dark grey surface), class 3 (light grey surface), and class 4 (medium grey surface). (F) The pointed-end particles were re-extracted and used as input to the supervised 3D classification, based on the unsupervised classification results, yielding class averages and associated frequencies marked in the same colors as references.

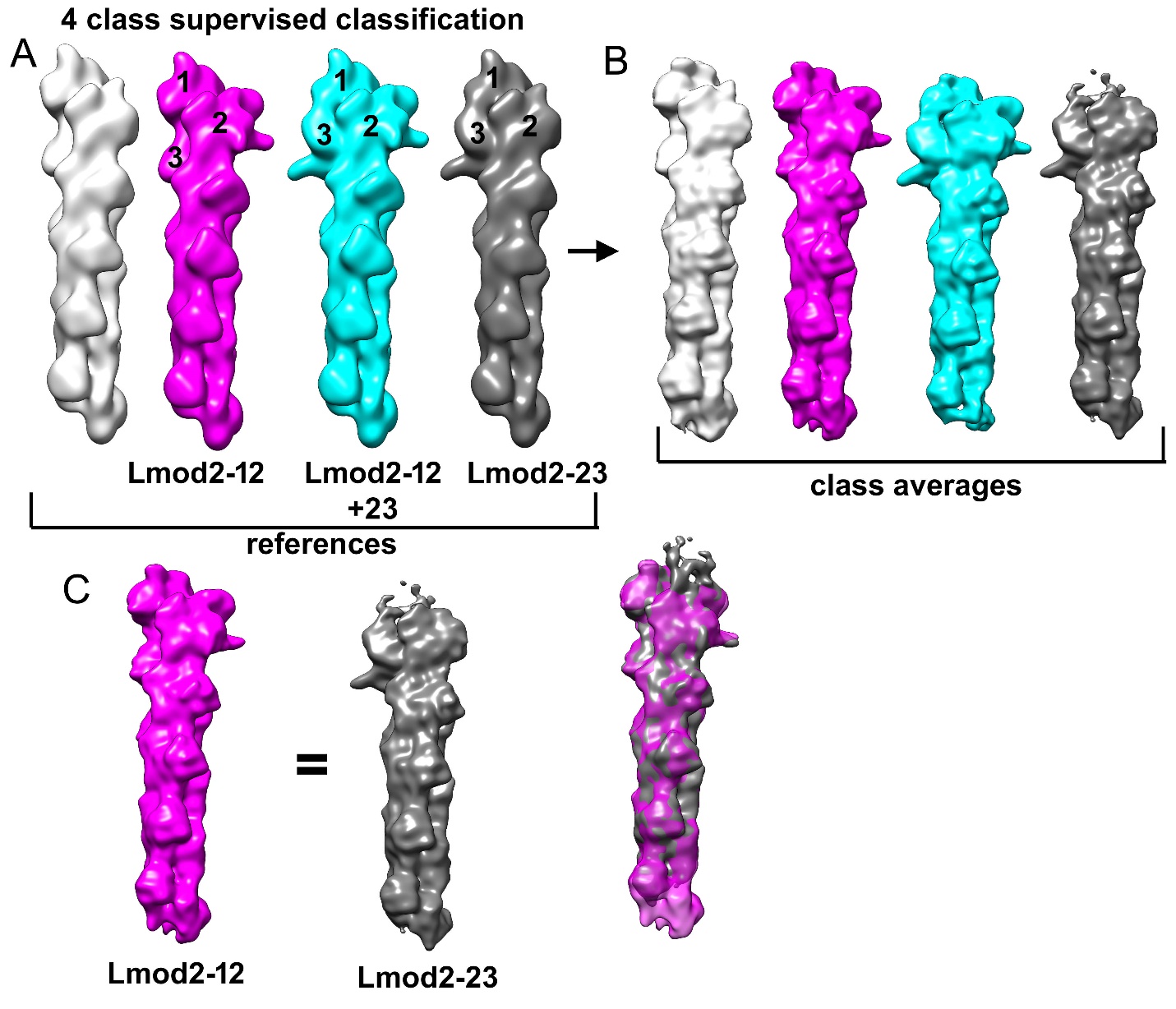

**Figure S3. 3D classification and verification of the pointed end structural modes.** (A) Four templates used in supervised classification of pointed ends are shown as follows: naked pointed end shown as a light grey surface, Lmod2 bound to actins 1 and 2 (i.e., Lmod2-12) shown as magenta surface, pointed end with Lmod2 bound to actins 1 and 2, and actins 2 and 3 (i.e., Lmod2-12+23) shown as a cyan surface, and pointed end possessing one Lmod2 bound to actins 2 and 3 (i.e., Lmod2-23) shown as dark grey surface. Actin protomers are numbered as in Fig 1(A-D). (B) Resultant class averages from 3D classification. (C) Lmod2-23 class average is equivalent to Lmod2-12.

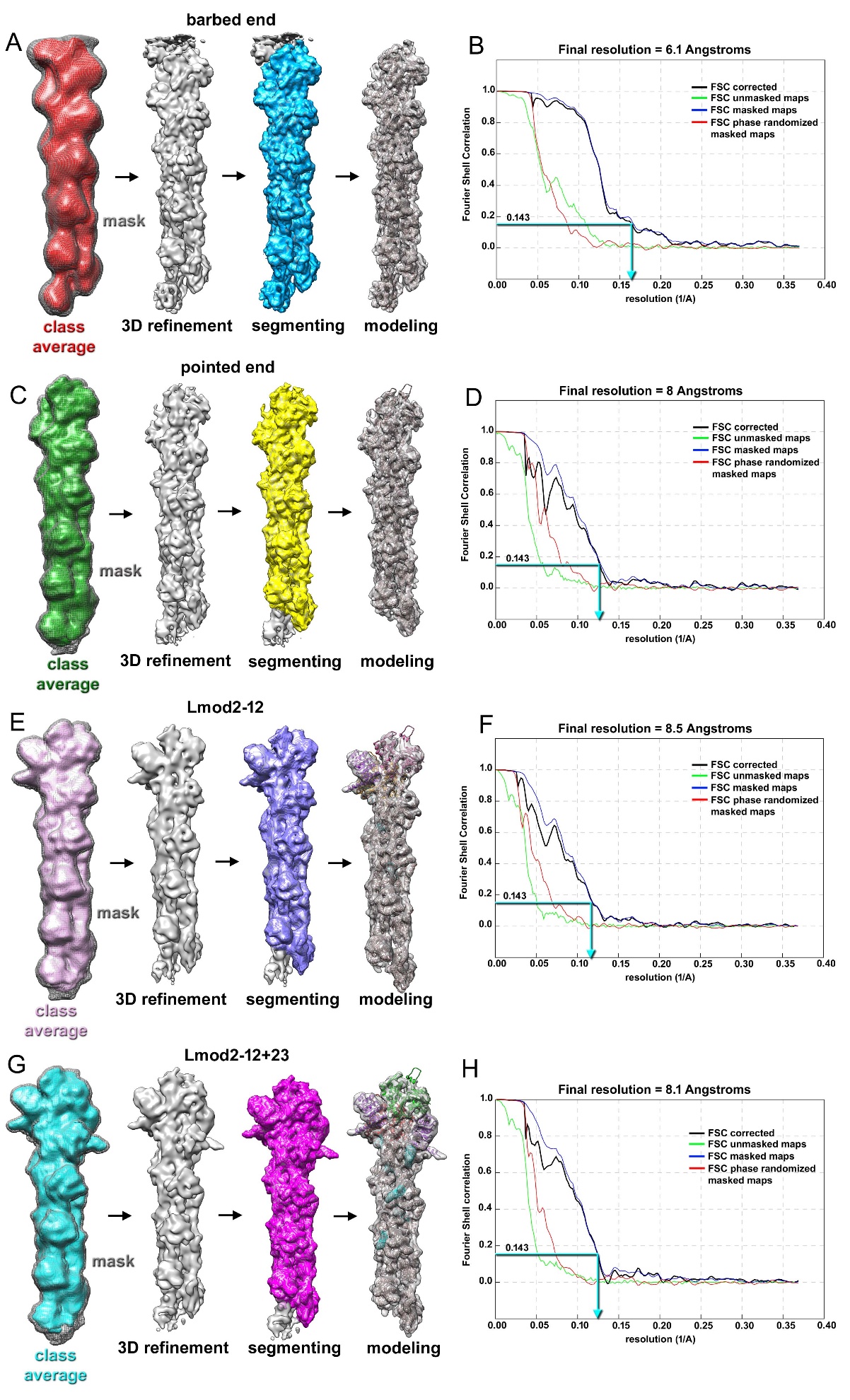

**Figure S4. 3D refinement and processing of structural classes.** (A) The class average for the barbed end served as the mask and reference (grey mesh and red surface, respectively) for 3D refinement. The region of interest was segmented from the 3D refinement and used for modeling. (B) Fourier shell correlation (FSC) plot. The map's global resolution was calculated using an FSC criterion of 0.143. (C-H) Pointed end, Lmod2-12, and Lmod2-12+23 class averages from the 3D sorting (green, pink, and cyan surfaces, respectively) were used as an initial reference and mask for 3D refinement, segmentation, and modeling (C, E, and G), and the global resolution (D, F, and H) was determined using an FSC criterion of 0.143. The processing is detailed in the Supplementary Information text.

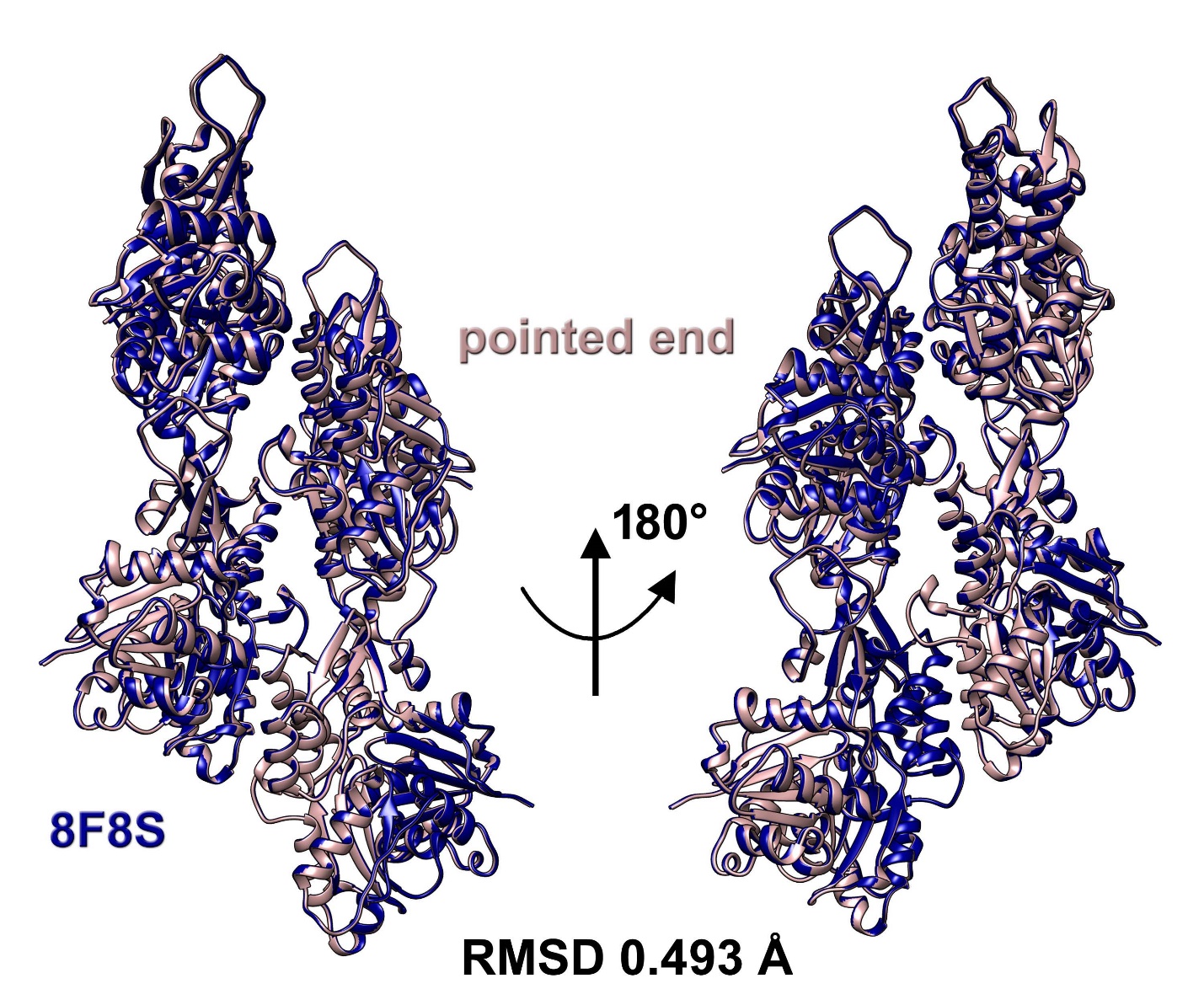

**Figure S5. Comparison of the pointed end model obtained in the current study with the previously published structure of skeletal F-actin pointed end.** The alignment of the naked pointed end model (rosy-brown ribbons) with the high-resolution atomic model of the skeletal F-actin (PDB 8F8S)^5^ (four actin protomers used) yielded an RMSD of 0.493 Å.

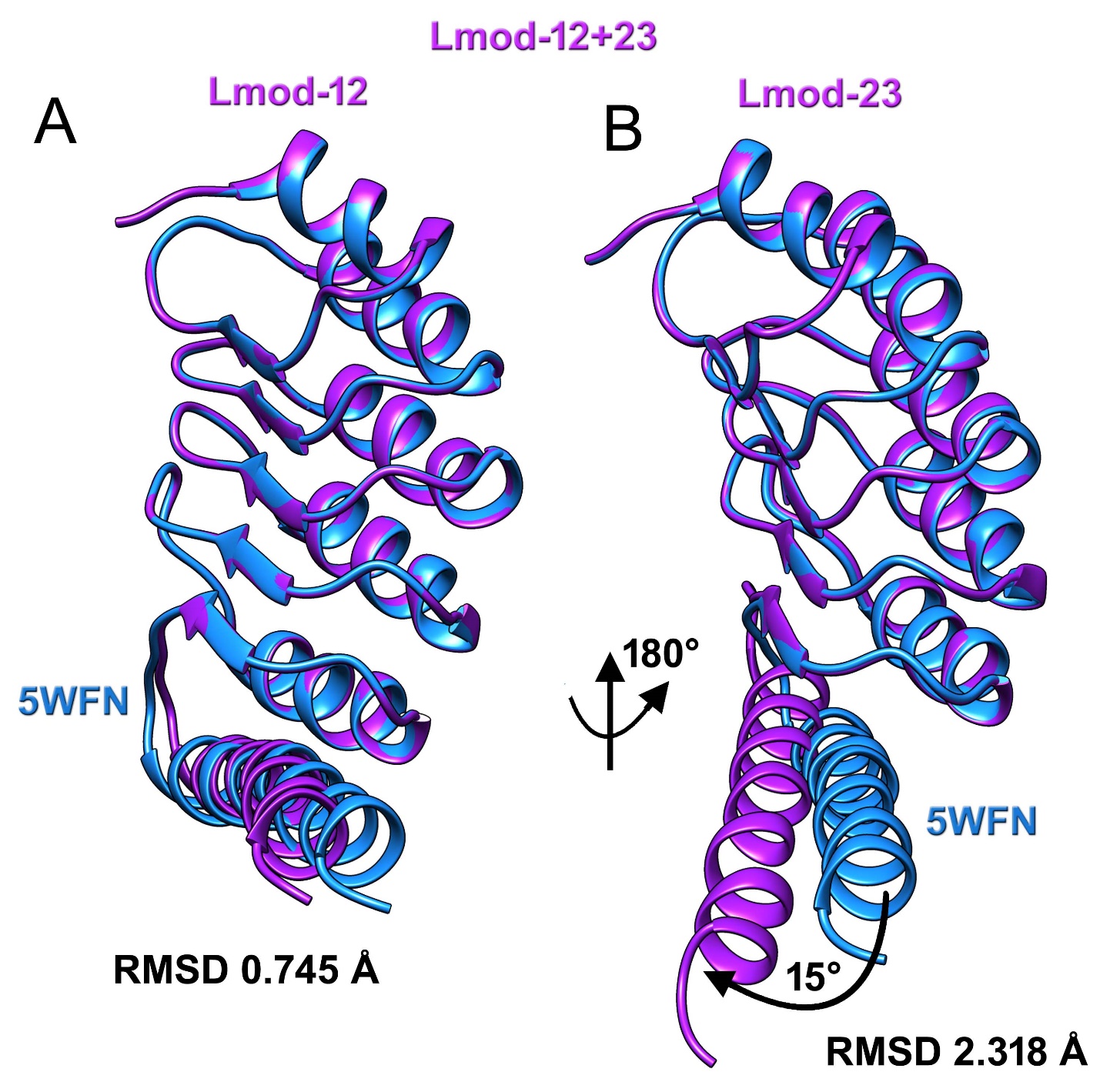

**Figure S6. Comparison of the LRR model obtained in the current study with the previously published crystal structure of the Lmod2-actin complex.** (A) LRR from the Lmod2-12+23 (purple ribbon) bound to actins 1 and 2 aligned with the crystal structure of the LRR (PDB 5WFN)^6^ (dodger blue ribbon). (B) LRR from the Lmod2-12+23 (purple ribbon) bound to actins 1 and 3 aligned with the crystal structure of the LRR (PDB 5WFN)^6^ (dodger blue ribbon). (A-B) Both LRRs in our atomic model exhibit excellent agreement with the crystal structure of Lmod2 bound to G-actin (PDB 5WFN), except for the C-terminal helix of LRR bound to actins 2 and 3 (B, black arrow), which has to be rotated by 15° to fit the density map.

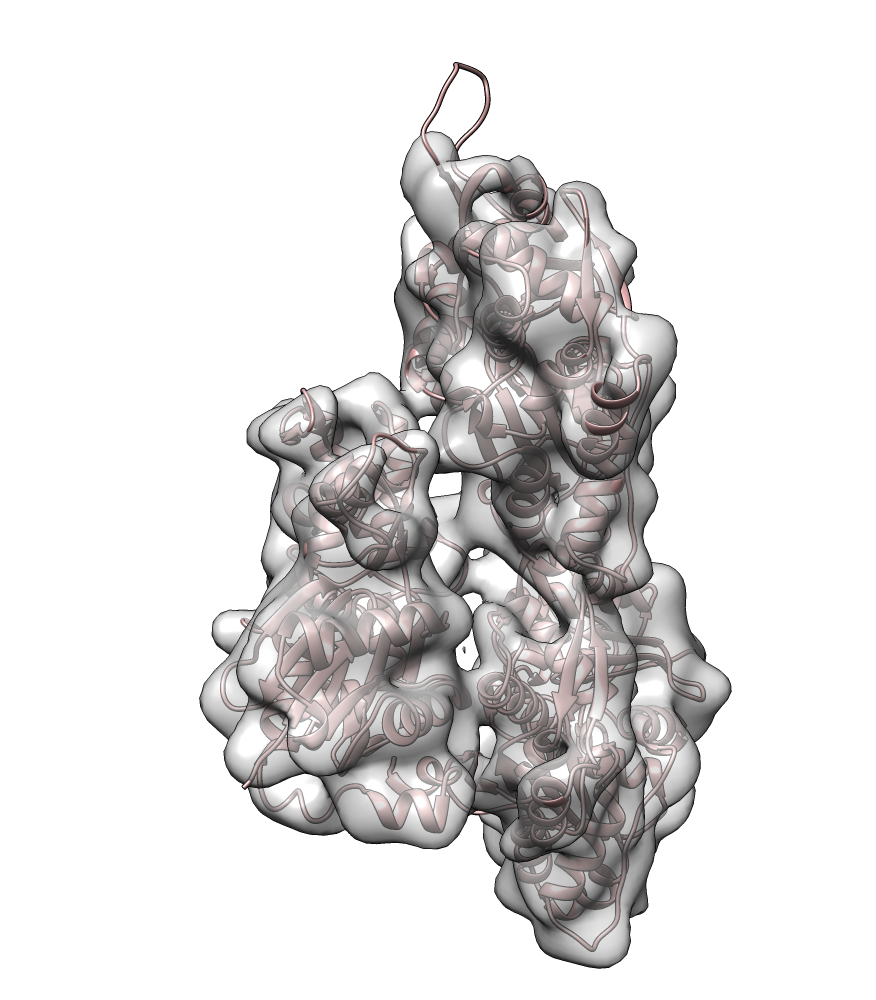

**Movie S1. Side view of the actin movement from the pointed end Lmod2-12 and Lmod2-12+23.** Cryo-EM models are shown as colored ribbons in the same color code as Figure 2. Cryo-EM maps are grey transparent surfaces.

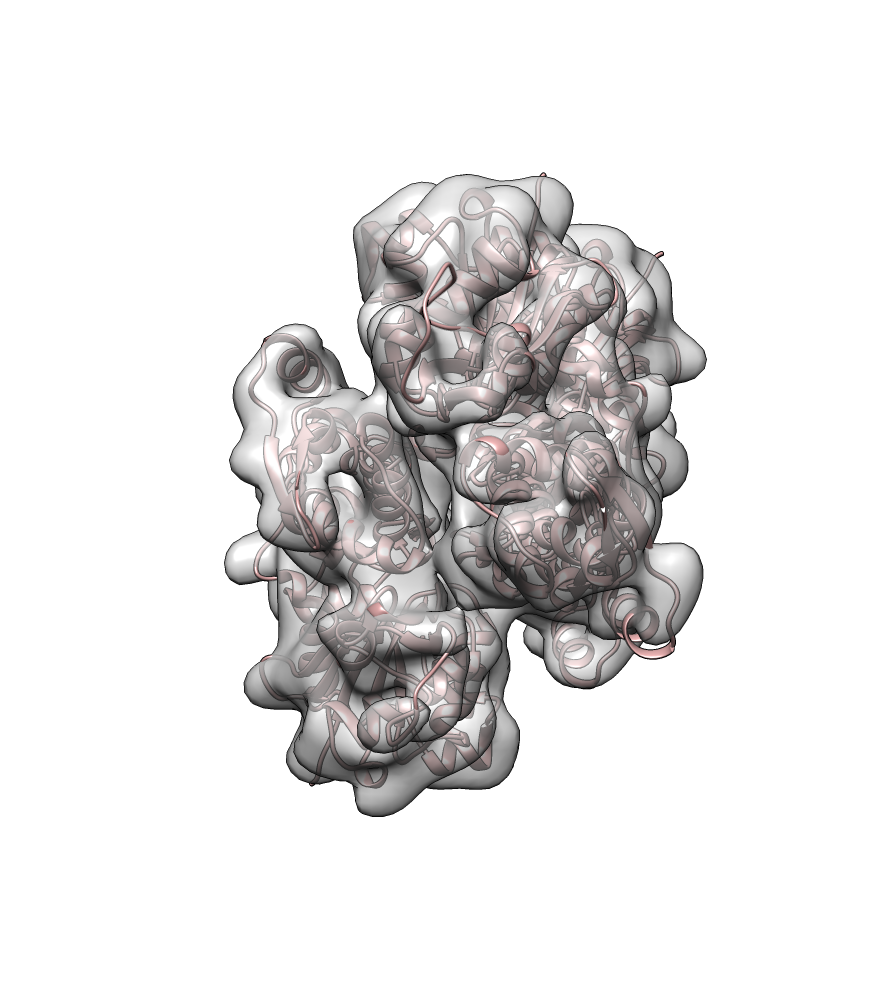

**Movie S2. Top view of the transition of the actin movement from the pointed end to Lmod2-12 and Lmod2-12+23.** Cryo-EM models are shown as colored ribbons in the same color code as Figure 2. Cryo-EM maps are grey transparent surfaces.

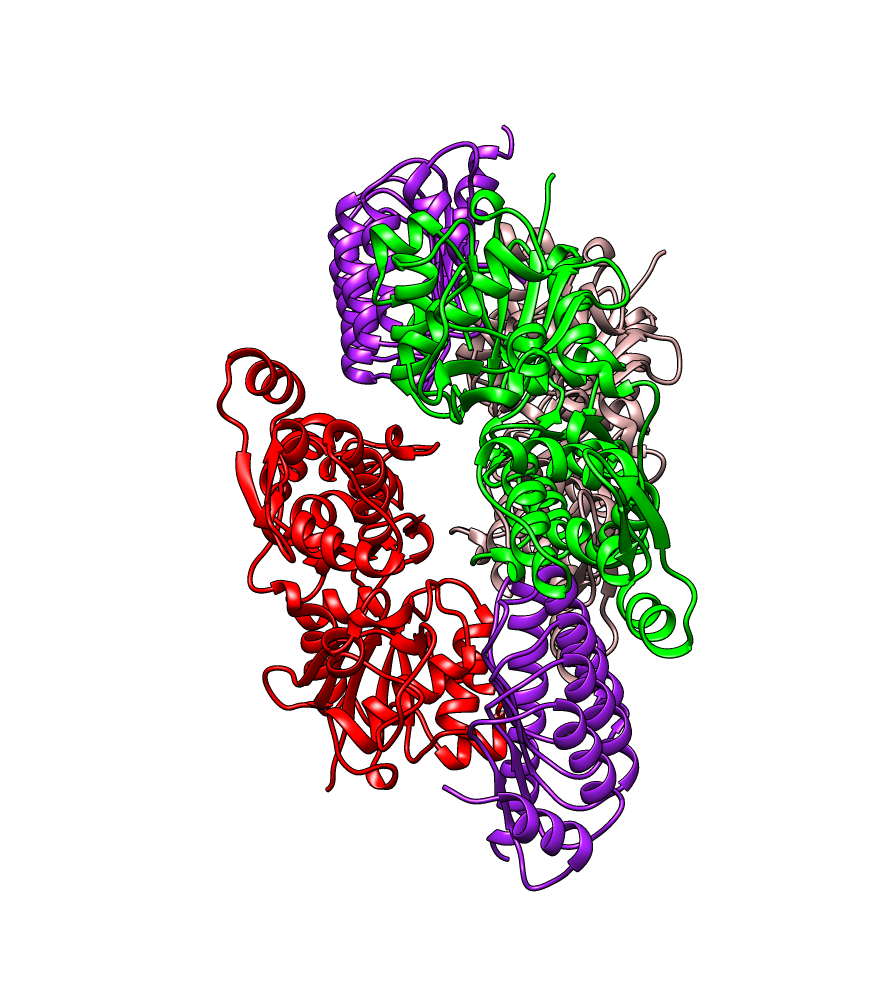

**Movie S3. Trajectory of the actin movement from the Lmod2-12+23 to Lmod2-12 to the pointed end.** Cryo-EM models are shown as colored ribbons in the same color code as Figure 2.

**Table S1.** Data collection and refinement statistics

| **Data collection and refinement statistics** | Barbed end | Naked pointed end | Pointed end in the presence of one Lmod2 | Pointed end in the presence of two Lmod2 |
| --- | --- | --- | --- | --- |
| **Data collection** |  |  |  |  |
| Magnification | 65,000 | 65,000 | 65,000 | 65,000 |
| Defocus range, µm | 0.5 – 3.5 | 0.5 – 3.5 | 0.5 – 3.5 | 0.5 – 3.5 |
| Voltage, kV | 300 | 300 | 300 | 300 |
| Microscope | Titan Krios | Titan Krios | Titan Krios | Titan Krios |
| Camera | K3 (super-resolution mode) | K3 (super-resolution mode) | K3 (super-resolution mode) | K3 (super-resolution mode) |
| Number of frames | 40 | 40 | 40 | 40 |
| Total electron dose, e^-^/Å^2^ | 34 | 34 | 34 | 34 |
| Frames used for final reconstruction and dose used for final reconstruction, e^-^/Å^2^ | Determined by Relion MotionCorr internal implementation | Determined by Relion MotionCorr internal implementation | Determined by Relion MotionCorr internal implementation | Determined by Relion MotionCorr internal implementation |
| Pixel size, Å/px | 0.678 | 0.678 | 0.678 | 0.678 |
| **Particle statistics** |  |  |  |  |
| Particles | 12,335 | 3,762 | 1,992 | 3,623 |
| Box size, Å | 540 | 540 | 540 | 540 |
| Pixel size, Å/px | 1.356 | 1.356 | 1.356 | 1.356 |
| **Resolution FSC 0.143, Å** | 6.1 Å | 8 Å | 8.5 Å | 8.1 Å |
| **Deposition ID**  **(EMD)** | 77128 | 77130 | 77131 | 77132 |

REFERENCES

1	Scheres, S. H. RELION: implementation of a Bayesian approach to cryo-EM structure determination. *J Struct Biol* **180**, 519-530 (2012). <https://doi.org/10.1016/j.jsb.2012.09.006>

2	Rohou, A. & Grigorieff, N. CTFFIND4: Fast and accurate defocus estimation from electron micrographs. *J Struct Biol* **192**, 216-221 (2015). <https://doi.org/10.1016/j.jsb.2015.08.008>

3	Pettersen, E. F. *et al.* UCSF Chimera--a visualization system for exploratory research and analysis. *J Comput Chem* **25**, 1605-1612 (2004). <https://doi.org/10.1002/jcc.20084>

4	Pintilie, G. D., Zhang, J., Goddard, T. D., Chiu, W. & Gossard, D. C. Quantitative analysis of cryo-EM density map segmentation by watershed and scale-space filtering, and fitting of structures by alignment to regions. *J Struct Biol* **170**, 427-438 (2010). <https://doi.org/10.1016/j.jsb.2010.03.007>

5	Carman, P. J., Barrie, K. R., Rebowski, G. & Dominguez, R. Structures of the free and capped ends of the actin filament. *Science* **380**, 1287-1292 (2023). <https://doi.org/10.1126/science.adg6812>

6	Boczkowska, M., Yurtsever, Z., Rebowski, G., Eck, M. J. & Dominguez, R. Crystal Structure of Leiomodin 2 in Complex with Actin: A Structural and Functional Reexamination. *Biophys J* **113**, 889-899 (2017). <https://doi.org/10.1016/j.bpj.2017.07.007>
