## Supplementary figures and images for "Structure of the Leiomodin-2 Regulated Actin Filament Pointed End Assembly from Profilactin"

### Supplementary movie 1

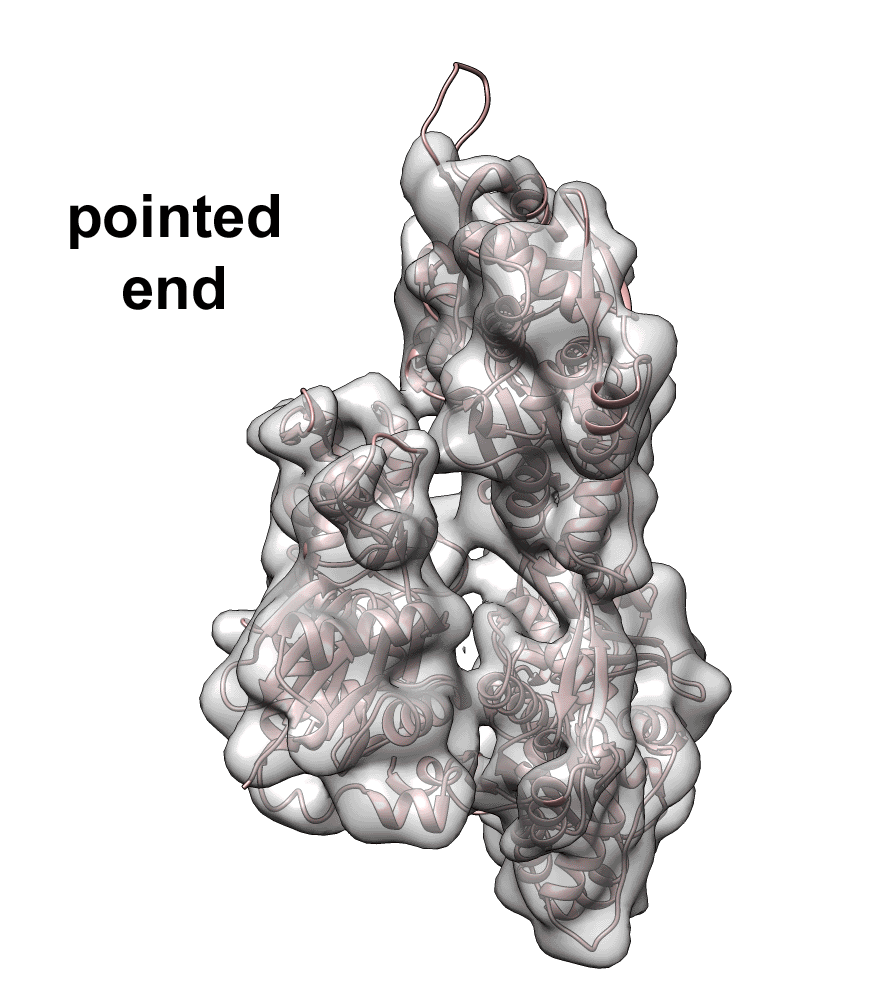

### Supplementary movie 2

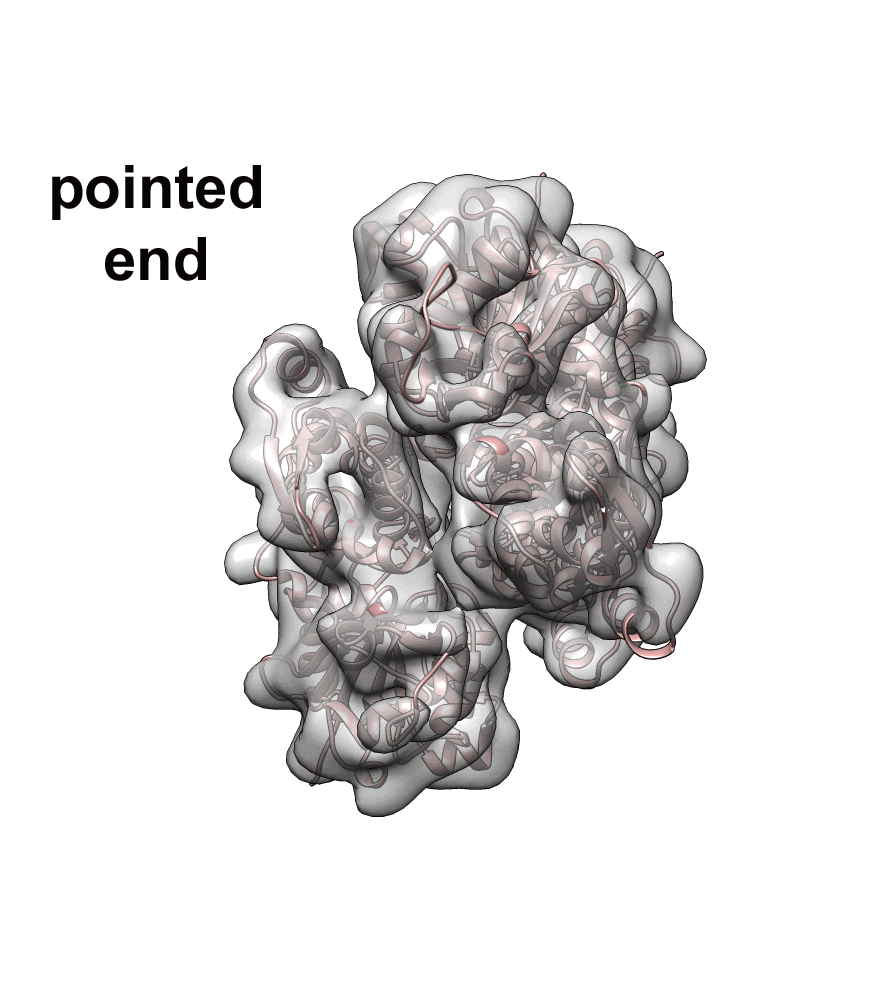

### Supplementary movie 3

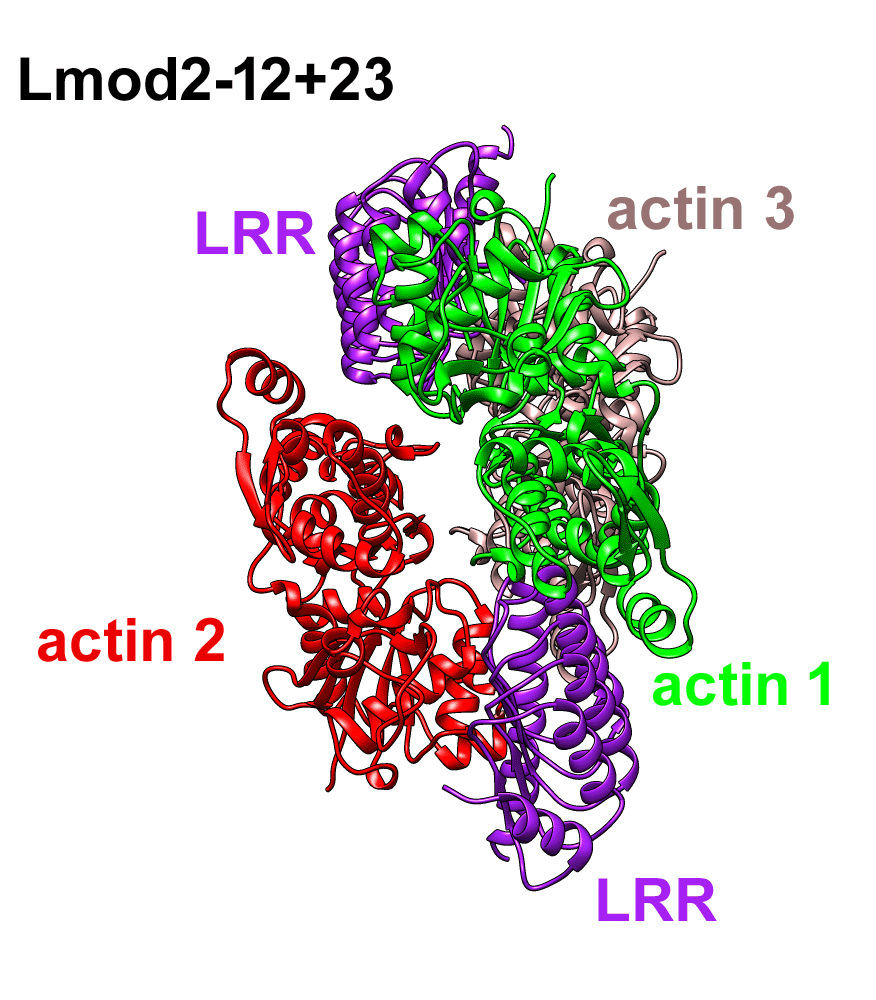
